# Improving selection efficiency in *C. americana* × *C. avellana* interspecific hybrids through the development of an indel-based genetic map

**DOI:** 10.1101/2023.02.05.527175

**Authors:** S.H. Brainard, J.A. Fischbach, L.C. Braun, J.C. Dawson

## Abstract

This study reports a genetic map created using a progeny family descended from the interspecific hazelnut cross *Corylus avellana* × *Corylus americana*. This research represents a critical step in the development of genomic tools that enable the deployment of next-generation sequencing methods in the breeding of hazelnut, specifically the improvement of well-adapted Midwestern hazelnut varieties. To produce this map, we first developed high-density molecular markers using short-read Illumina sequencing of genotype-by-sequencing libraries. By aligning reads to a newly assembled reference genome for *C. americana*, we were able to identify over 75,000 high-quality indel-based polymorphisms across an F_1_ experimental population. These markers exhibited both high allele depth coverage, and low linkage disequilibrium, making them well-suited to genetic map development. We constructed such a map using 95 individuals from a single F1 family, demonstrating the utility of next-generation sequencing to efficiently and accurately generate high-density genetic maps. This research will improve the efficiency of breeding efforts, both through the validation of specific molecular markers that are associated with agronomically-relevant traits in breeding populations of interest.

## INTRODUCTION

Hazelnuts (*Corylus* spp.) are a globally significant nut crop, with annual production of over 1 million tons, being produced across 34 countries (FAOSTAT, 2022). Its present cultivation is, however, limited to areas with climatic conditions approximating the Mediterranean region in which the dominant cultivars of *C. avellana* were bred. This significantly restricts expansion of U.S. production beyond the Willamette Valley in Oregon, where over 95% of current U.S. acreage is located. In this context, the shrub species *C. americana*, native to much of the eastern U.S., offers a valuable pool of germplasm for breeding more widely-adapted hazelnut varieties, which can exhibit vigorous growth in cooler environments, and robust resistance to aggressive pathogens such as the diverse, endemic fungal disease Eastern Filbert Blight (*Anisogramma anomala*).

Developing genetic tools is critical in this regard, both to identify QTL for marker-assisted selection, but also to efficiently generate molecular markers for use in genomic prediction. Genetic maps are an indispensable tool, in particular with regards to the former goal. While numerous genetic maps have been published using SSR (Mehlenbacher et al., 2006;

Mehlenbacher and Bhattarai, 2018) and SNP markers (Rowley et al., 2018) in *C. avellana* backgrounds, to date, the only maps which have been generated for *C. avellana* × *C. americana* hybrid hazelnuts have utilized wild *C. americana* selections (Lombardoni et al., 2022). This study addresses this knowledge gap through the construction of a genetic map using an F_1_ progeny family descended from a cross between Oregon and Midwestern hazelnut varieties ‘Jefferson’ and ‘Eric4-21’, respectively. In addition, this map employs a marker technology not previously used in map construction in hazelnuts: multiallelic indel markers identified from Illumina short reads. This study therefore not only extends genetic mapping to a novel interspecific hybrid population, but also employs a novel and powerful marker type. By combining the cost-efficiency of next-generation sequencing, with the additional information regarding recombination events contained in markers with more than two allelic states, the map presented here is both high-density, and extremely accurate.

## MATERIALS AND METHODS

### Sample preparation and sequencing

An F_1_ population of 95 individuals was used to construct the genetic map presented here. The maternal parent was a variety developed by the Upper Midwest Hazelnut Development Initiative, *C. americana* × *C. avellana* ‘Eric4-21’. The paternal parent was the commercially important variety bred by Oregon State University, *C. avellana* ‘Jefferson’. This cross was made at the University of Minnesota in 2015, and replicated in 2016 with the progeny being planted at a University of Minnesota research station in Rosemount, MN in 2016 and 2017. Leaf tissue from the parental lines and F_1_ progeny was obtained immediately following leaf-out in the spring of 2020. Tissue was lyophilized, and genomic DNA was extracted and purified at the University of Wisconsin Biotechnology Center. Reduced-representation sequencing libraries were prepared using a double digestion with the restriction enzymes *Nsi*I and *Bfa*I following the methodology described by Elshire et al., 2011. Illumina GBS barcodes and adapters were ligated to DNA fragments, and paired-end reads (2 × 150 bp) were generated using an Illumina NovaSeq 6000, generating an average of 12 million reads per sample, or roughly 10x coverage. This represented 772,351 unique 64mers (i.e., 49.43 Mb of unique sequence, representing 9.57% of the 473 Mb carrot genome)

### Genetic map construction

Following trimming and demultiplexing of Illumina reads (performed using the custom Java software gbsTools; https://github.com/shbrainard/gbsTools), phased, haplotype-based markers were called using Stacks 2 (Rochette et al., 2019). The parameter ‘gt-alpha’ was set to 0.01 to increase filtering stringency, and confidence in identified genotypes. Markers were called using a reference genome assembly for *C. americana* accession ‘Winkler’ (PI 557019, NCGR, Corvallis, OR). Markers were then filtered using bcftools (Danecek et al., 2021) to remove sites in high linkage disequilibrium with each other (R^2^ > 0.95), leaving 78,079 markers (an average of ∼7,000 markers per chromosome). Markers were additionally selected on the basis of total allelic variants segregating in the population, filtering for segregation types A1, A2, and B3.7 (following the notation of (Wu et al., 2002)), leaving only markers with either three or four alleles.

Maps were then constructed using the R package onemap (Margarido et al., 2007). In brief, markers were first filtered for missing data by removing sites with no genotype calls across more than 5% of all individuals. Two-point recombination frequencies were then calculated for all markers, and all four possible phase configurations using maximum likelihood estimation. Because only those markers with fully informative segregation types were selected, the correct phase between any pair of markers was able to be determined on the basis of maximum likelihood estimations themselves. Hierarchical clustering was then performed to form linkage groups, and a Hidden Markov Model (HMM) was used to simultaneously order and phase markers within these groups. A global error rate of 0.05 was used in the HMM, which together with removal of unlinked markers controlled for inflation. The Kosambi mapping function was used to convert recombination frequencies to genetic distances.

#### Data availability

The genetic map is available for download and use via Figshare at https://doi.org/10.6084/m9.figshare.21902193.

## RESULTS AND DISCUSSION

The distribution of markers included in the final genetic map across chromosomes and segregation types are shown in Table 1. A key method for evaluating genetic map quality are heatmaps, which visualize the matrix of all pairwise recombination frequencies, given a specified marker ordering and phasing. A correct ordering would exhibit minimum recombination frequencies between adjacent pairs of markers (the super- and sub-diagonals of the matrices), with frequencies increasing monotonically with distance between markers (reaching a maximum in the upper left and lower right corners of the matrix, which represent recombination between markers at opposite ends of a given linkage group). Representative heatmaps for three of the eleven linkage groups in hazelnut are shown in Fig. 1. These heatmaps illustrate the successful ordering of markers within linkage groups using the onemap pipeline. Furthermore, they demonstrate the value of utilizing multiallelic markers: all cells in the heatmap have a color, a consequence of the fact that recombination is observable in both parents for all pairs of markers, and thus maximum likelihood estimates of recombination frequency can be obtained for each pair. This is not the case, for example, when using SNP markers which typically exhibit only 2 allelic states. Markers with two alleles are not fully informative, given the 4 haplotypes segregating in an F_1_ family descended from outcrossed parents. While SSR markers can also provide multiallelic resolution in assessing polymorphisms, the cost associated with genotyping SSR’s typically limits the size of genetic maps to only a few hundred, while indels obtained for GBS data can be genotyped for a fraction of the cost, allowing the use of the thousands of markers included in this study.

**Table 1.** Number of markers per chromosome which were included in the final genetic map, as well as the corresponding distribution of these markers across the three segregation types (A.1, A.2 and B3.7).

| **Chromosome** | **# of markers** | **A.1** | **A.2** | **B3.7** |
| --- | --- | --- | --- | --- |
| 1 | 208 | 43 | 110 | 55 |
| 2 | 201 | 47 | 119 | 35 |
| 3 | 134 | 35 | 86 | 13 |
| 4 | 106 | 28 | 59 | 19 |
| 5 | 129 | 30 | 78 | 21 |
| 6 | 89 | 23 | 56 | 10 |
| 7 | 430 | 8 | 138 | 284 |
| 8 | 93 | 20 | 62 | 11 |
| 9 | 81 | 23 | 52 | 6 |
| 10 | 146 | 30 | 68 | 48 |
| 11 | 74 | 17 | 45 | 12 |

**Figure 1.**
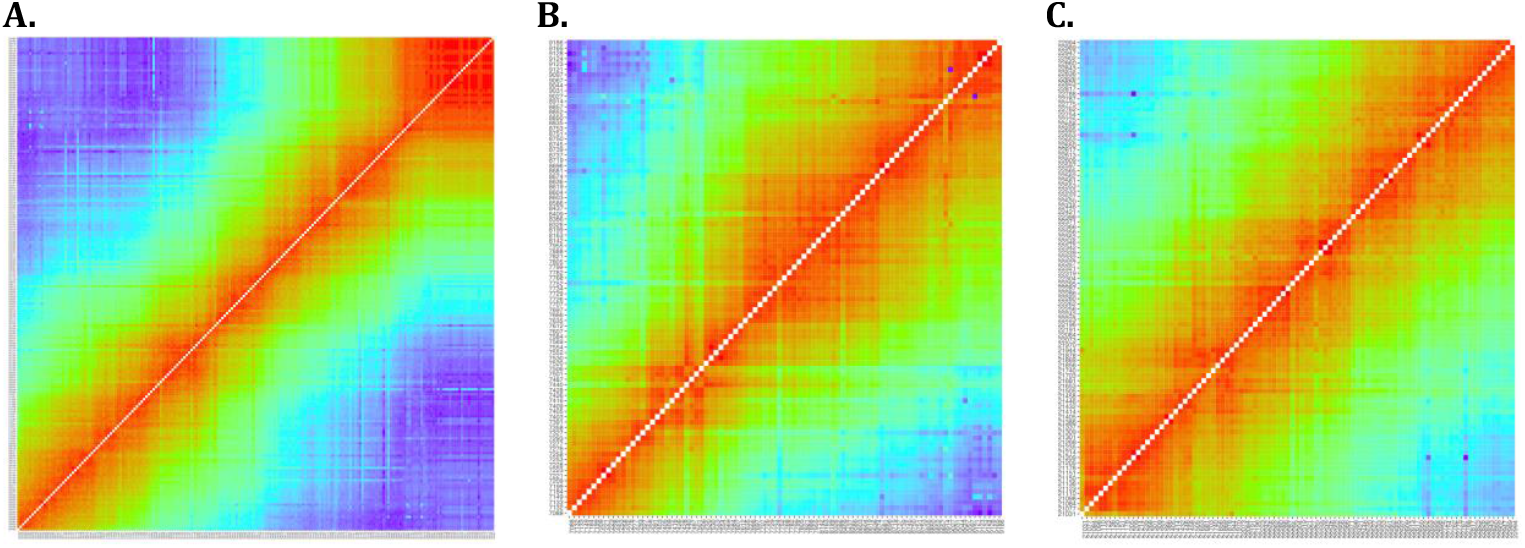
Heatmaps of recombination frequencies generated by the R package onemap for linkage groups corresponding to chromosomes 2 (A), 6 (B) and 8 (C). The color spectrum ranges from red (indicating a recombination frequency of 0) to blue (indicating a recombination frequency of 0.5).

**Figure 2.**
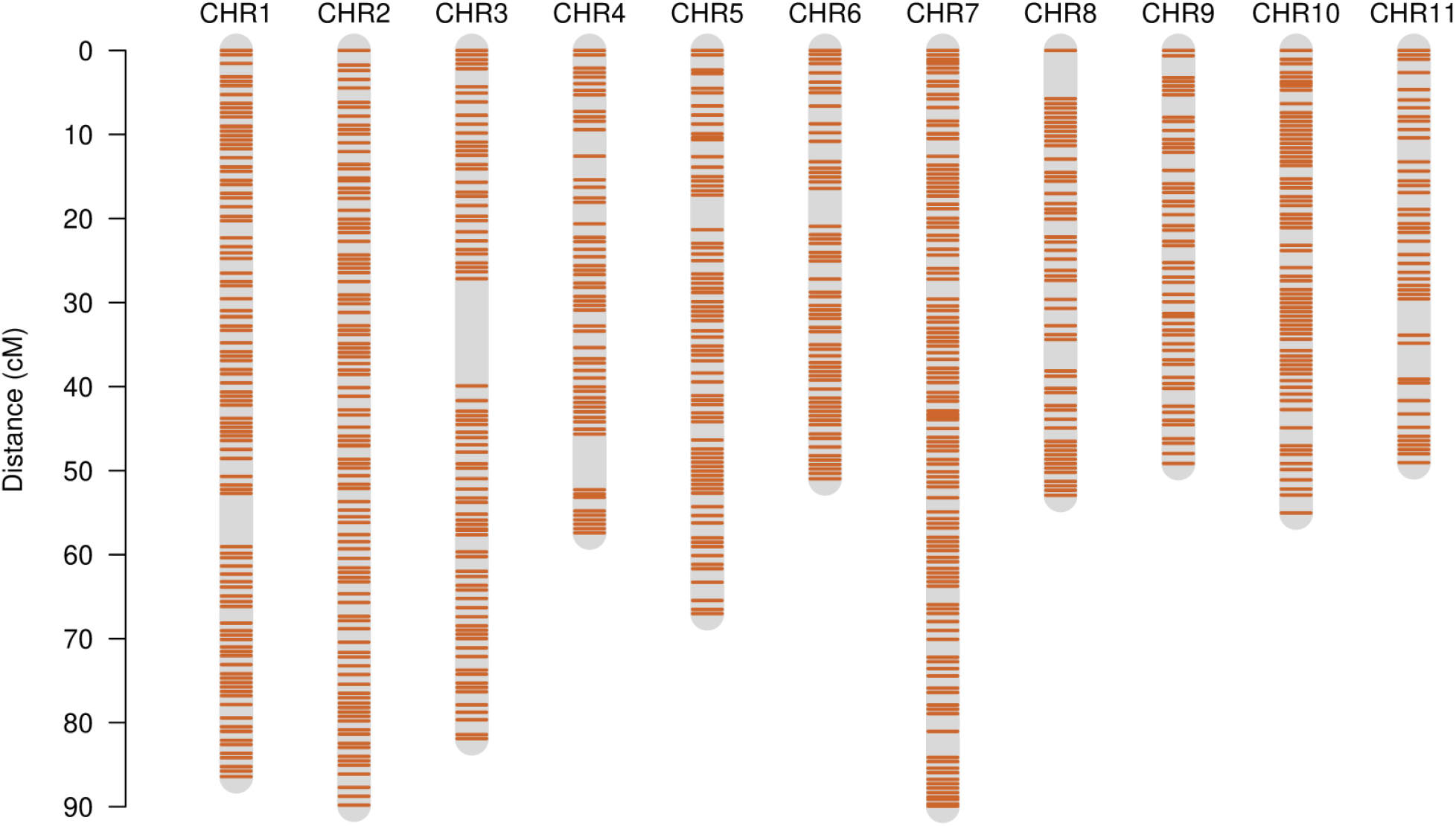
The distribution of markers along the genetic map for the F_1_ family Eric4-21 × Jefferson. Genetic distance is expressed in cM while linkage group IDs have been replaced with their respective chromosome number from the reference assembly used to identify polymorphisms.

A second key measure of genetic map quality is the estimated length of the map. Given the relatively small size of hazelnut chromosomes (with an average physical size of 29.78 Mb), under normal biological assumptions and a standard mapping function, a given linkage group should not be longer than 100 cM. Inflated map lengths in excess of this upper bound are nevertheless frequently observed by not controlling for genotyping errors, which increase estimates of recombination frequency. Using multi-point regression when converting genetic distances to the final genetic map counteracts this tendency. The use of multiallelic markers in the onemap pipeline was shown to produce a high-quality map also in this regard: linkage groups are no longer than 89.9 cM, with an average length of 66.3 cM, and a total length of 729.6 cM.

## CONCLUSION

This study presents a genetic map constructed using an F_1_ progeny family descended from a Midwest-adapted hybrid hazelnut variety, and a commercially significant Oregon State release. The use of multi-allelic markers was demonstrated to facilitate the development of a very high-quality map, exhibiting accurate ordering and phasing of maternal and paternal haplotypes within the F_1_ family, with no apparent inflation. This map adds to the growing body of such studies using multiallelic markers to improve map quality, and eventually statistic power in subsequent QTL analyses (Casas et al., 2018; Thérèse Navarro et al., 2022). As such, it provides researchers with a valuable resource in investigating the genetic control of agronomically-relevant traits, and will hopefully accelerate the expansion of the cultivated range of hazelnut varieties.

## ACKNOWLEDGEMENTS

We thank Mark Hamann of the University of Minnesota, and Shawn Mehlenbacher of Oregon State University for their assistance in providing leaf tissue samples from F_1_ progeny and parental varieties. The University of Wisconsin Bioinformatics Resource Center provided support in running the Stacks 2 pipeline. Funding for this research was provided by the USDA-NIFA SCRI Grant No. H007913501.

